# Interplay between a bacterial pathogen and its host in rainbow trout isogenic lines with contrasted susceptibility to Cold Water Disease

**DOI:** 10.1101/2023.02.23.529686

**Authors:** Bo-Hyung Lee, Edwige Quillet, Dimitri Rigaudeau, Nicolas Dechamp, Eric Duchaud, Jean-François Bernardet, Pierre Boudinot, Tatiana Rochat

## Abstract

Infectious diseases are a major constraint on aquaculture. Genetic lines with different susceptibilities to diseases are useful models to identify resistance mechanisms to pathogens and to improve prophylaxis. Bacterial cold-water disease (BCWD) caused by *Flavobacterium psychrophilum* represents a major threat for freshwater salmonid farming worldwide. A collection of rainbow trout (*Oncorhynchus mykiss*) isogenic lines was previously produced from a French domestic population. Here, we compared BCWD resistance phenotypes using a subset of isogenic lines chosen for their contrasted susceptibilities to *F. psychrophilum*. We applied individual monitoring to document the infection process, including time-course quantification of bacteremia and innate immune response. Strikingly, BCWD resistance was correlated with a lower bacterial growth rate in blood. Several immune genes were expressed at higher levels in resistant fish regardless of infection: the Type II arginase (*arg2*), a marker for M2 macrophages involved in anti-inflammatory responses and tissue repair, and two Toll-like receptors (*tlr2*/*tlr7*), responsible for pathogen detection and inflammatory responses. This study highlights the importance of innate and intrinsic defense mechanisms in determining the outcome of *F. psychrophilum* infections, and illustrates that non-lethal time-course blood sampling for individual monitoring of bacteremia is a powerful tool to resolve within-host pathogen behavior in bacterial fish diseases.

## 1. Introduction

Aquaculture has been the world’s fastest-growing food production over the last four decades and the “blue revolution” is considered to be one of the most promising part of the global food system to meet the demand of quality proteins for increasing population [1]. Salmonid fish production is one example for such aquaculture expansion. Among many factors of concern including production efficiency and durability, fish health and welfare are of high importance and infectious diseases represent the main bottleneck for sustainable development of the aquaculture industry. Bacterial cold-water disease (BCWD), also known as rainbow trout fry syndrome, is a devastating bacterial disease in freshwater salmonid farming worldwide. *Flavobacterium psychrophilum*, the causative agent, affects high commercial value salmonid species such as rainbow trout (*Oncorhynchus mykiss*) and Atlantic salmon (*Salmo salar*). The treatment of BCWD exclusively relies on antibiotics administration in fish feed, which bears a risk of emergence of antibiotic resistance in fisheries and surrounding environmental microbiota.

The general symptoms of BCWD are well documented. Infected fish present signs of tissues erosion and necrotic lesions, typically in the caudal fin and the jaw, skin ulcers, exophthalmia, as well as inflammation in organs [2]. In rainbow trout fry, that are particularly susceptible, the disease results in haemorrhagic septicaemia and high mortality. Cellular and molecular mechanisms behind pathogenesis are still unclear. Several factors influencing the outcome of infection have been reported in the past decades based on rainbow trout experimental infection studies, including alterations of skin integrity, environmental stresses, fish immune status and genetic background of both *F. psychrophilum* and its host [3–7].

Genetic resistance to *F. psychrophilum* infections has been established with moderate heritability estimated in domestic rainbow trout broodstocks [8–10]. Selective breeding allows genetic gain and appears as an attractive strategy to reduce the impacts of outbreaks [11–13]. Rainbow trout resistance to BCWD has been investigated in resistant/susceptible lines using linkage and genome-wide association studies. Characterization of experimental ARS-Fp-R and ARS-Fp-S lines obtained by the USDA through family-based selection and breeding populations identified multiple Quantitative Trait Loci (QTLs) of BCWD resistance, and genomic positions were recently localized more precisely for two of them, providing promising information on putative mechanisms underlying BCWD resistance [14–18]. At INRAE, a collection of doubled-haploid isogenic rainbow trout lines was screened for resistance to various diseases including several viruses and pathogenic bacteria, some of which showed contrasted susceptibility to *F. psychrophilum* [19,20]. BCWD resistance phenotyping of a double haploid QTL family produced by a cross between two isogenic lines, AP2 (Resistant) and B57 (Susceptible), using injection and immersion methods identified 15 QTLs associated to resistance traits, 12 of which were specific to the infection route [5]. These susceptible isogenic lines are also an attractive resource to achieve standardized and reliable experimental BCWD models [21–23]. In addition to QTLs discovery, comparative transcriptomic studies of resistant and susceptible rainbow trout lines also suggested relevant immune genes that could be responsible for resistance to BCWD [24,25].

Assessment of disease resistance in fish was mainly quantified by recording mortality as a binary parameter as well as time-to-death values. Non-lethal biopsies are a routine practice in many models to monitor parameters such as pathogen loads or antibody titers at individual host level; however, their application in aquatic animals has not been obvious. Recently non-lethal repeated blood collections was successfully implemented in Atlantic salmon [26].

Here, we report a comparative study of BCWD resistance phenotypes for a collection of rainbow trout isogenic lines that show contrasted susceptibilities to *F. psychrophilum* infection. For the first time in the investigation of bacterial infection in rainbow trout, we applied individual monitoring to document the infection process using time-course determination of bacteremia and host immune responses in blood.

## 2. Materials and methods

### 2.1. Bacterial strain and culture condition

The virulent *F. psychrophilum* strain FRGDSA 1882/11 isolated in France from diseased rainbow trout and belonging to the clonal complex CC-ST90 (ST108) and the serotype Th was used [27,28]. Bacterial cultures were carried out in tryptone yeast extract salts broth (TYES) at 200 rpm and 18°C as previously described (Zhu et al 2022). Late-exponential phase cultures (OD600 of 1.0 ± 0.1, equivalent to 10^9^ CFU/ml) were used for infection challenges. Bacterial concentration was determined by colony forming unit (CFU) counting after plating on TYES agar supplemented with 5% fetal calf serum (FCS) as previously described (Zhu et al 2022).

### 2.2. Rainbow trout experimental infections

Rainbow trout isogenic lines originated from the INRAE reference synthetic line (Sy) [29]. Gynogenesis was performed during two generations to produce all female and homozygous doubled haploid lines. Each isogenic line was further propagated by single pair mating between a female and a sex-reversed male as previously described [19]. Fish were reared at 10°C in recirculating aquaculture system in 30-liter tanks with dechlorinated and UV treated tap water, then transferred to a biosafety level 2 (BSL2) zone in 15-liter tanks with flow water (1 renewal per hour) for infection experiments. Fish were fasted for 24 h or 48 h prior to infection by injection or immersion, respectively. Five experimental infections were performed (Table 1); they differed in the infection route (intramuscular injection *versus* bath) and the size of fish (in fry, juvenile or adult stage).

**Table 1.** *F. psychrophilum* experimental infection schemes

| Trial | Isogenic line | Condition | Infection route | Infectious doses (CFU)<sup>a</sup> |  | Age (month) | Average weight (g) | Total number of fish |
| --- | --- | --- | --- | --- | --- | --- | --- | --- |
| I | A03 | *F. psychrophilum* | Bath | 8.9 x 10<sup>6</sup> | 7.4 x 10<sup>6</sup> | 4 | 1.2 | 100 |
|  | G17 |  |  | 8.2 x 10<sup>6</sup> | 1.0 x 10<sup>7</sup> | 5 | 2.0 | 100 |
|  | A36 |  |  | 6.4 x 10<sup>6</sup> | 1.4 x 10<sup>7</sup> | 4.5 | 1.3 | 100 |
|  | B57 |  |  | 9.9 x 10<sup>6</sup> | 9.0 x 10<sup>6</sup> | 4.5 | 1.0 | 100 |
|  | AP2 |  |  | 5.4 x 10<sup>6</sup> | 8.5 x 10<sup>6</sup> | 4 | 1.1 | 100 |
| II | A03 | *F. psychrophilum* | IM injection | 3.0 x 10<sup>2</sup> | 4.5 x 10<sup>2</sup> | 8 | 9.9 | 91 |
|  | G17 |  |  |  |  | 9 | 9.6 | 89 |
|  | A36 |  |  |  |  | 8.5 | 10.0 | 87 |
|  | B57 |  |  |  |  | 8.5 | 12.2 | 85 |
|  | AP2 |  |  |  |  | 8 | 10.3 | 93 |
| III | A36 | *F. psychrophilum* | IM injection | 2.4 x 10<sup>4</sup> |  | 13 | 61.1 | 26 |
|  | B57 |  |  |  |  | 13 | 101.9 | 26 |
|  | AP2 |  |  |  |  | 14 | 98.4 | 26 |
| IV | A36 | *F. psychrophilum* | IM injection | 2.1 x 10<sup>4</sup> |  | 17 | 149.7 | 26 |
|  | B57 |  |  |  |  | 17 | 303.7 | 26 |
|  | AP2 |  |  |  |  | 18 | 376.6 | 26 |
|  | A36 | TYES control | IM injection | - |  | 17 | 131.6 | 10 |
|  | B57 |  |  |  |  | 17 | 320.0 | 10 |
|  | AP2 |  |  |  |  | 18 | 347.0 | 10 |
| V | A36 | *F. psychrophilum* | IM injection | 1.1 x 10<sup>4</sup> |  | 27 | 398.5 | 4 |
|  | B57 |  |  |  |  | 27 | 767.8 | 4 |
|  | AP2 |  |  |  |  | 28 | 969.2 | 4 |

<sup>a</sup> For Trial I, bacterial concentration is indicated as CFU/mL of water for each duplicate tank at the beginning of immersion. For Trial II-V, infectious doses correspond to colony forming units (CFU) injected per fish. Two independent cultures were used for Trial II duplicated tanks.

Five isogenic lines namely A03, A36, AP2, B57 and G17, were challenged via immersion and injection routes. For the bath challenge (Trial I), 100 fish of each isogenic line were distributed in duplicate tanks at the fry stage (mean body weight 1.33 ± 0.37 g). For each duplicate tank, a biological independent bacterial culture was 100-fold diluted into 10 l of tank water. Bacteria were maintained in contact with fish for 18 h by stopping the water flow and were subsequently removed by restoring the flow. During the immersion, water was maintained at 10°C under vigorous aeration ([O_2_] > 9 mg l^-1^). *F. psychrophilum* bacterial counts were determined at the beginning of the immersion challenge by plating serial dilutions of water sample on TYES FCS agar. The IM injection challenge (Trial II) was carried out on 445 fish that were digitally tagged (Tiny puce, Biolog-id) and distributed in two replicates in two 300-liter tanks (mean number of fish: 44 per isogenic line and per tank). Bacterial suspensions (one per duplicate tank) were prepared by 5 × 10^5^-fold dilution of two independent cultures. Fish were anesthetized in 100 mg l^-1^ tricaine (MS222, Sigma) and 50 µl of bacterial suspension were delivered in the dorsal muscle close to the dorsal fin. Dead fish were removed from aquaria twice a day during 25 days to monitor the survival kinetics.

Three IM injection trials (III-V; Table 1) were performed at adult stage using the isogenic lines A36, AP2 and B57. Bacterial suspensions were prepared by 10^4^-fold dilution of bacterial cultures and 100 µl were used for IM injection as described above. Control groups fish were injected with 100 µl of sterile TYES. In trial III and IV, groups of 26 fish were randomly distributed into two compartments of a 300-liter tank separated using a perforated polyvinyl chloride divider to establish the survival kinetic (n=10) and sampling (n=16) for each isogenic line. In trial III, blood sampling was performed for *F. psychrophilum* cells enumeration at 3 days post infection (dpi, n=6) and at the onset of mortality (n=6; corresponding to 7 dpi for A36 and B57 isogenic lines and 10 dpi for AP2). In trial IV, 16 fish in total of the sampling compartment were digitally tagged and divided in 2 subgroups using fin clipping method (to avoid unnecessary anesthesia) for individual monitoring assay and alternative blood collection. Fish were anaesthetized and 150 µl of blood was rapidly collected in a heparinized syringe from the caudal vein for bacterial enumeration. In trial IV, bacterial loads were determined in the spleen of dead fish from the survival monitoring compartment: for A36 and B57, 5 spleens were sampled from dead fish; for AP2, spleens were sampled from 2 dead fish and from 5 fish euthanized at 19 dpi to respect humane endpoints (severe skin lesions and persistent morbidity signs). In trial V, 12 fish (4 per isogenic line) reared in the same 300-liter tank were marked by digital tagging and used for individual monitoring of gene expression. Blood was collected as described above at 3 time points: before infection (0 dpi); early-stage infection (3 dpi); moribund stage (corresponding to 8, 11 and 14 dpi for A36, B57 and AP2, respectively). Blood was immediately processed for RNA extraction.

Statistical differences were analysed with GraphPad Prism 8.2.0.

### 2.3. RNA extraction and qRT-PCR

Sixty µl of blood was transferred to a 1.4 mm ceramic beads pre-filled tube containing 600 µL of RLT from the RNeasy Mini Kit (Qiagen), samples were homogenized, RNA was extracted and then treated using RNase-Free DNase Set following the manufacturer’s instructions. qPCR was performed on CFX system following manufacturer’s instruction (BIO-RAD) using primers listed in Table 2. Mean Ct values were calculated using technical triplicates and normalized by geometric mean of two reference genes (ELF1α and -actin).

**Table 2.**
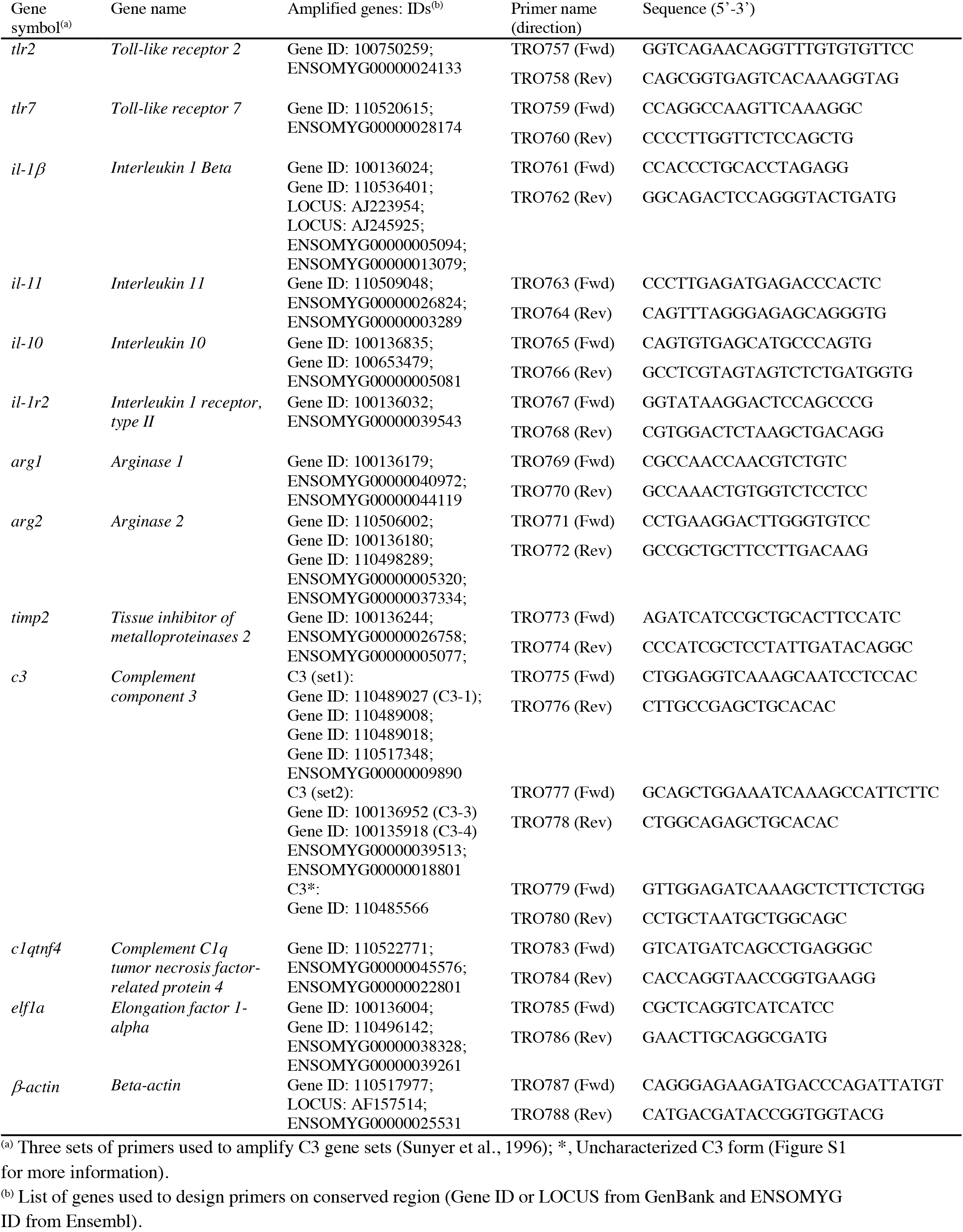
Primer sequences used in the qRT-PCR analysis

### 2.4. Ethics statements

Animal experiments and sampling were performed at the INRAE-IERP fish facilities of Jouy-en-Josas (France) in accordance with the European Directive 2010/2063/UE. All animal work was approved by the Direction of the Veterinary Services of Versailles, France (building agreement number C78-720) and by the ethics committee of the INRAE Center in Jouy-en-Josas (COMETHEA n°45), France (authorization numbers #12//53 and #2015100215242446).

## 2. Results

### 2.1. Susceptibility to F. psychrophilum infection differs between rainbow trout isogenic lines

The BCWD resistance/susceptibility of five rainbow trout isogenic lines was compared using experimental infection at the fry stage (Table 1). Waterborne infection resulted in a large range in the degree of susceptibility with final survival proportions ranging from 0% to 94% (Fig. 1A). While lines AP2 and A03 were highly resistant to a *F. psychrophilum* immersion challenge, A36 showed rapid death that reached 95 % mortality at 16 dpi. B57 and G17 showed intermediate mortality with 51 % and 38 % at the end of the observation time (dpi 28), respectively. Based on Kaplan-Meier curves, differences were significant between the following phenotypic groups: Resistant (AP2, A03), Intermediate (B57, G17) and Susceptible (A36).

**Fig. 1.**
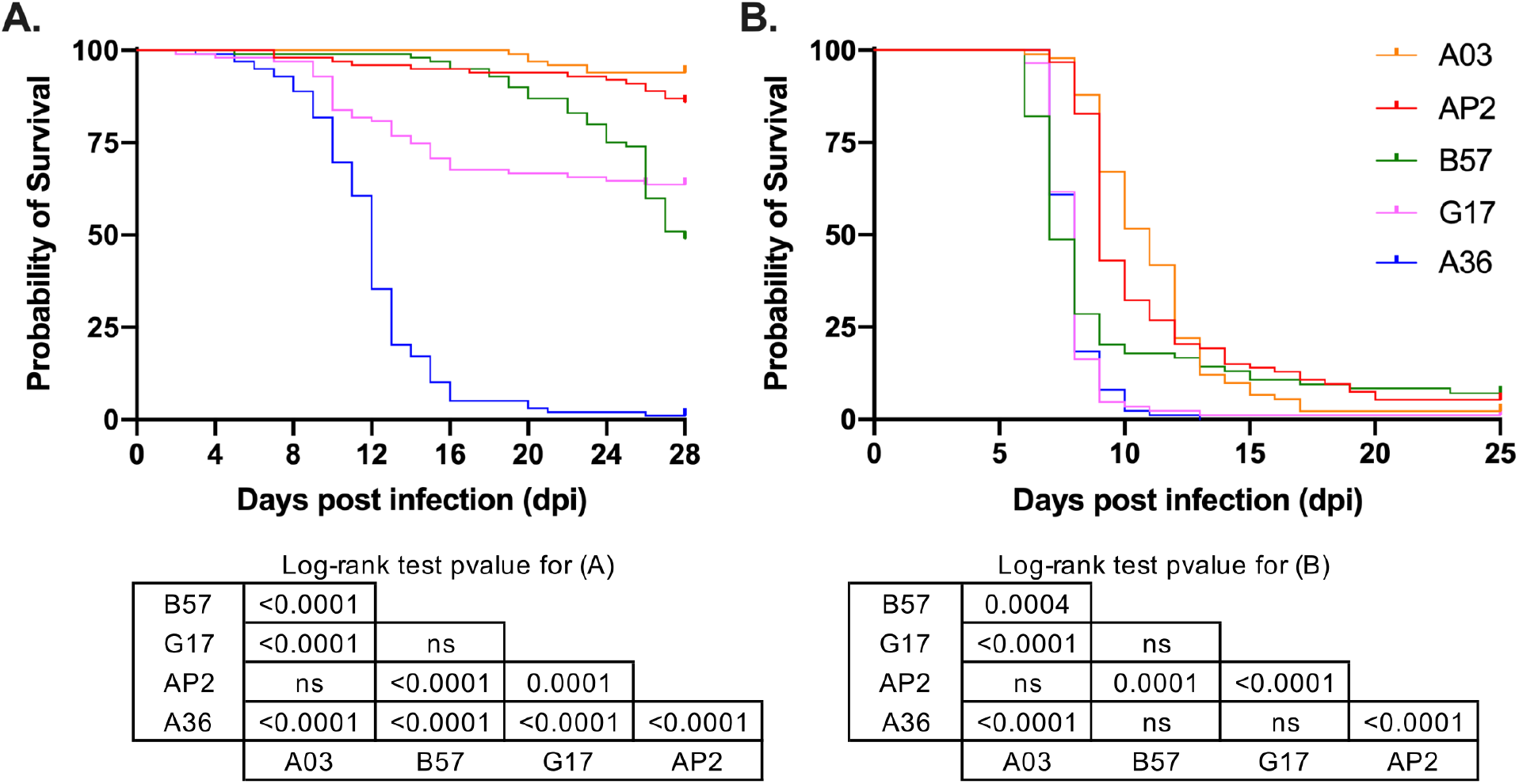
Rainbow trout isogenic lines reveal contrasted susceptibilities to *F. psychrophilum* infection. Kaplan-Meier survival curves of rainbow trout fry infected via immersion (A) and IM injection (B) show significant differences among some isogenic lines. Lower panel: tables with Mantel-Cox log-rank test p-values for bath (left) and IM injection (right) infection.

In contrast to the immersion challenge, all injected groups reached > 94 % mortality at the end of the observation (25 dpi) after intramuscular injection despite a low infectious dose (3.8 ±0.7 × 10^2^ CFU) (Fig. 1B). This result is in line with previous studies pointing out the important role of mucosal and skin barriers in BCWD resistance [3,4,30] Time-to-death variations resulted in significant differences between the survival curves of isogenic lines. Lines of the Resistant group (AP2 and A03) showed moderate decline in the survival curves, while those of the Susceptible group (A36 and G17) revealed rapid mortality at 7-9 dpi. The survival curve of line B57 was not significantly different from those of lines A36 and G17, although it was positioned between the survival curves of the two phenotypic groups due to extended survival of B57 fish which were still alive after 9 dpi (20%). The B57 phenotype was therefore classified as “Intermediate”. Most importantly, the ranking of the five isogenic lines established for their BCWD susceptibility was conserved between immersion and injection.

Based on the range of BCWD susceptibility observed at the fry stage, three isogenic lines were selected for further investigations, namely A36, B57, and AP2 as representative of Susceptible, Intermediate and Resistant phenotypic groups, respectively. Additional IM injection trials (Table 1) were performed in order to establish their respective BCWD resistance phenotype at adult stage (Trial III).

Significant differences were observed between the survival kinetics of adult fish: mortality started from 7 dpi for lines A36 and B57 and from 10 dpi for AP2 (Fig. 2A). While the survival curve of A36 displayed a steep decline resulting in the death of all individuals within two days with brief appearance of morbidity signs, those of B57 and AP2 displayed a gradual decline as time passed, reaching 90 % cumulative mortality at 13 dpi. Bacterial loads in blood were determined by sampling 6 fish per group after euthanasia at 3 dpi and at the onset of mortality, a time point that differed between host genotypes (Fig. 2B). Bacteremia levels were similar at the early stage of infection and increased in all isogenic lines at late stage. Strikingly, it reached significantly higher levels in Susceptible line A36 compared to Resistant line AP2, with Intermediate B57 values being positioned in between. These results suggest that within-host bacterial growth rate and host response to infection differ between isogenic lines resulting in different outcomes of BCWD infection. To test this hypothesis, experimental infection was reproduced using individual monitoring method (Table 1) to record simultaneously the survival curves and the time-course bacterial loads in blood (Trial IV) and to determine immune responses kinetics (Trial V).

**Fig. 2.**
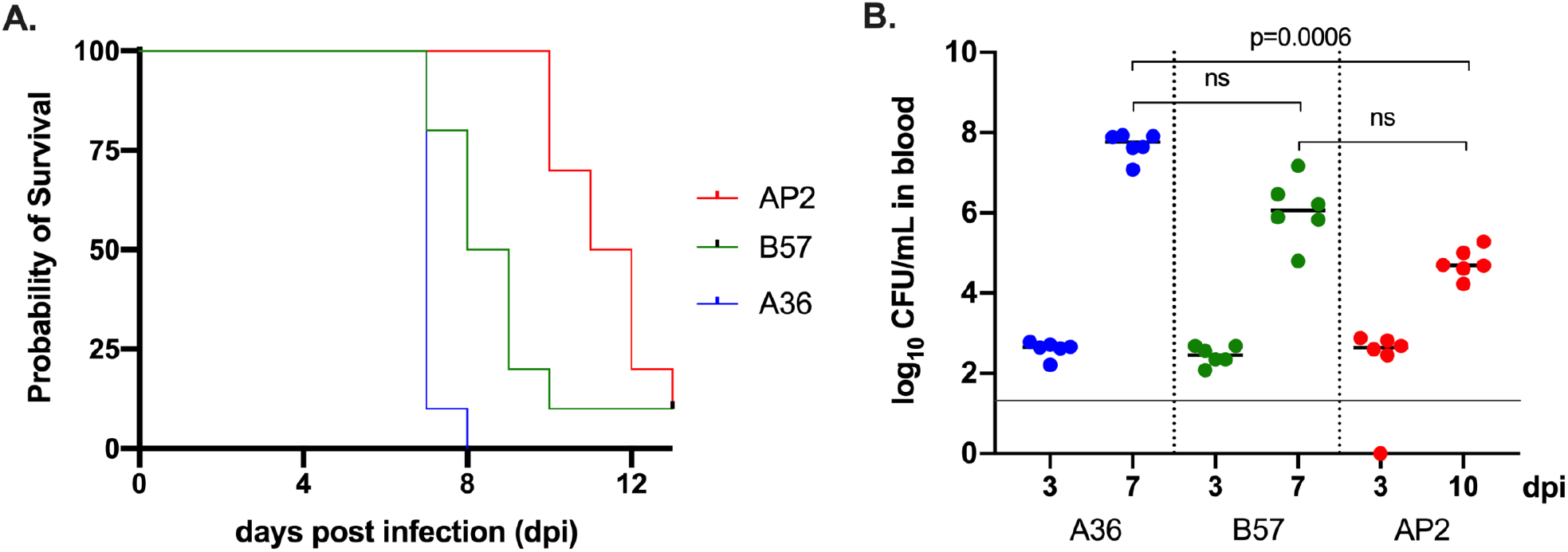
BCWD resistance phenotypes of three rainbow trout isogenic lines following infection by injection at the adult stage. A. Kaplan-Meier survival curves of fish infected via IM injection (Table 1, Trial III). Significant differences were observed between all survival curves (Mantel-Cox log-rank test p-values = 0.0184 between B57 and AP2; = 0.0011 between B57 and A36; <0.0001 between AP2 and A36). B. Bacterial loads in the blood were determined at 3 dpi (n=6) and at the onset of mortality (n=6; 7 dpi for A36 and B57; 10 dpi for AP2). Significant differences were observed between A36 and AP2 isogenic lines at late stage (Kruskal-Wallis test p-values).

### 2.2. BCWD resistance is associated with within-host bacterial growth control

An optimized sampling plan for individual monitoring of bacteremia was designed in a way to minimize putative effects of recurrent blood collection. Survival was recorded with unsampled individuals in a second tank compartment to follow the course of infection (Fig. 3A).

**Fig. 3.**
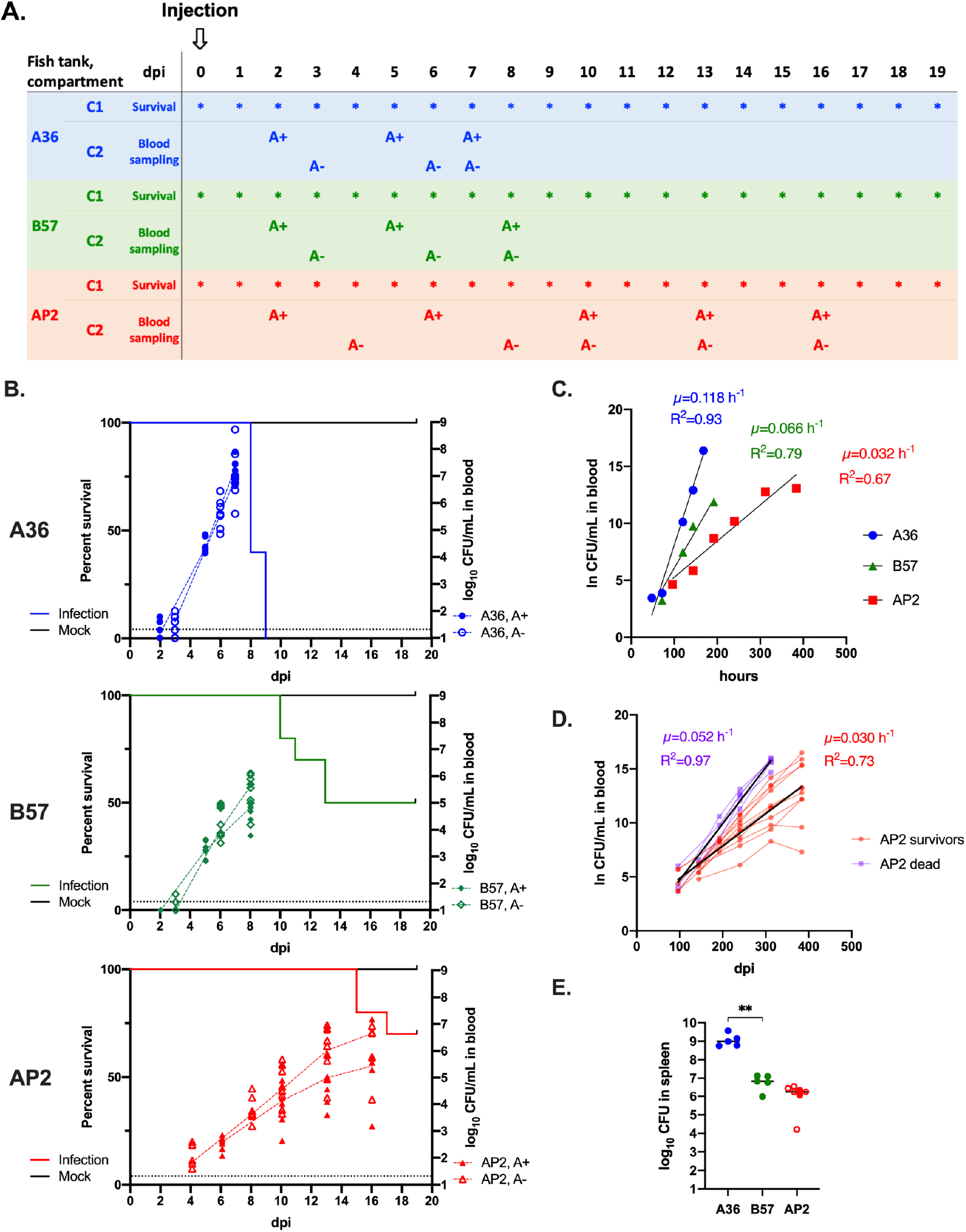
Survival kinetics of three isogenic lines correlate with bacterial load in blood in adult fish during BCWD experimental infection by injection. A. Plan of Trial IV (Table 1). A total of 26 fish were infected and distributed into two compartments (C1/C2) of a tank for each isogenic line: one group for survival monitoring and spleen sampling of dead fish (C1; n=10) and the other one for successive non-lethal blood samplings (C2; n=16). In the latter compartment, fish were divided into two sub-groups of 8 fish by clipping of the adipose fin (A+, intact adipose fin; A-, clipped adipose fin) for alternative sampling during the course of infection every 3 days and at the moribund stage (7 dpi for A36; 8 dpi for B57 and 10, 13, and 16 dpi for AP2). Five AP2 fish died on 15 dpi and blood samples were collected from the survivors on 16 dpi. B. Kaplan-Meier estimation of survival (left axis; plain line) and level of bacteremia expressed in log_10_ CFU.ml^-1^ (right axis). Dashed lines connect medians of each sub-group (filled symbols: A+; open symbols: A-). C. Curves fitted using time-course of ln-transformed CFU.ml^-1^ values from 16 fish per isogenic line using the linear regression function of the GraphPad Prism 8 software. Pairwise comparisons showed different slopes (p-value 0.0001). µ: growth rate (h^-1^); R^2^: coefficient of determination. D. Individual kinetics of bacteremia in two AP2 sub-groups ‘dead’ (violet: 5 fish dead at 15 dpi) and ‘survivors’ (red: 11 fish alive at 16 dpi). E. Bacterial loads in spleen of dead fish (n=2; filled symbols) or fish euthanized at 19 dpi after persistent morbidity signs (n=5; open symbols). ** Mann-Whitney test p-value < 0.01.

Differences between survival curves of the three isogenic lines observed in Trial III were reproduced with the same ranking. However, we observed a delay in the beginning of death events and a lower mortality rate for AP2 and B57 (Fig. 3B; Table 1). AP2 fish demonstrated behavioral symptoms including lethargy and swim defects from 10 dpi, however endured and survived better to the infection compared to Trial III. At 19 dpi, AP2 fish developed massive viscous cystic nodules at the injection site and were euthanized as they reached humane endpoint criteria. In the blood sampling compartments, mortality events tend to be more frequent and to occur earlier compared to unsampled fish in the survival compartments: all B57 fish were dead at 13 dpi; 31% of AP2 fish died at 15 dpi and the remaining survivors were euthanized at 16 dpi after the last blood sampling. Additional stresses induced by successive handling and sampling may have worsened the disease.

The level of bacteremia correlated with BCWD susceptibility determined using survival estimates as a proxy (Fig. 3B) and the results were consistent with those measured in Trial III (Fig. 2B). At the moribund stage, A36 fish demonstrated the highest bacterial burden in blood by reaching log_10_ CFU ml^-1^ of 7.1 ± 0.7 (mean, SD) at 7 dpi while B57 reached log_10_ CFU ml^-1^ of 5.2 ± 0.7 at 8 dpi. During the time when AP2 fish showed behavioral signs while resisting to infection, the average bacteremia levels measured before death events (at 13 dpi) and in survivors (at 16 dpi) were comparable (log_10_ CFU ml^-1^ of 5.5 ± 1.1 and 5.7 ± 1.2, respectively), but high inter-individual variations were observed (Fig. 3B, lower panel). Within-host *F. psychrophilum* growth rate (*µ*) in each isogenic line was then estimated by linear regression using the ln-transformed CFU.ml^-1^ values in blood as a function of time and bacterial generation time was deduced (Fig. 3C). Strikingly, bacterial generation time was strongly dependent of the host genotype, with significant differences between A36 (5.9 hours), B57 (10.5 hours) and AP2 (21.8 hours). For AP2, average bacteremia levels apparently reached a plateau at 13 dpi (Fig. 3B). We questioned whether this observation reflected an effective bacterial growth arrest from 13 dpi or resulted from average calculation. When considering individual time-course bacteremia data for the two AP2 sub-groups ‘dead’ (5 fish dead at 15 dpi) and ‘survivors’ (11 fish alive at 16 dpi), the bacterial load was higher in blood for all fish of the ‘dead’ group compared to the ‘survivors’ and the respective bacterial growth rates were significantly different (Fig. 3D; p=0.0001). Individual curves of bacterial loads in AP2 blood clearly showed a significant inter-individual variation of pathogen growth.

Bacterial loads in the spleen upon death followed the same trend as in blood: bacterial load reached very high levels in Susceptible line A36 (median value of 1.0 × 10^9^ CFU) compared to Intermediate line B57 (median value of 6.9 × 10^6^ CFU) (Fig. 3E). For AP2, bacterial loads were determined for dead and euthanized fish: values were intertwined and bacterial loads were similar to those observed for B57 (median value, 1.8 × 10^6^ CFU). Altogether, these results indicate that BCWD resistance in rainbow trout is associated with a better control of pathogen growth.

### 2.3. Differential expression of immune genes in isogenic lines upon F. psychrophilum infection

Putative differences in the establishment of effective immunological control were investigated further in the three isogenic lines using individual monitoring of the expression of relevant immune genes in blood (Trial V). The relative expression of 11 immune genes/gene families (Table 2; Fig. S1) selected based on previous *F. psychrophilum* infection studies [24,25] was determined before infection (0 dpi), at early stage (3 dpi) and in moribund with A36 at 0 dpi used as a reference condition, and significant differences were observed for 8 genes (Fig. 4; Fig. S2). Generally, induced genes showed highest expression at the early stage of infection confirming the establishment of infection and immune reactions. Moreover, significant differences in basal expression levels of some genes were observed among the isogenic lines.

**Fig. 4.**
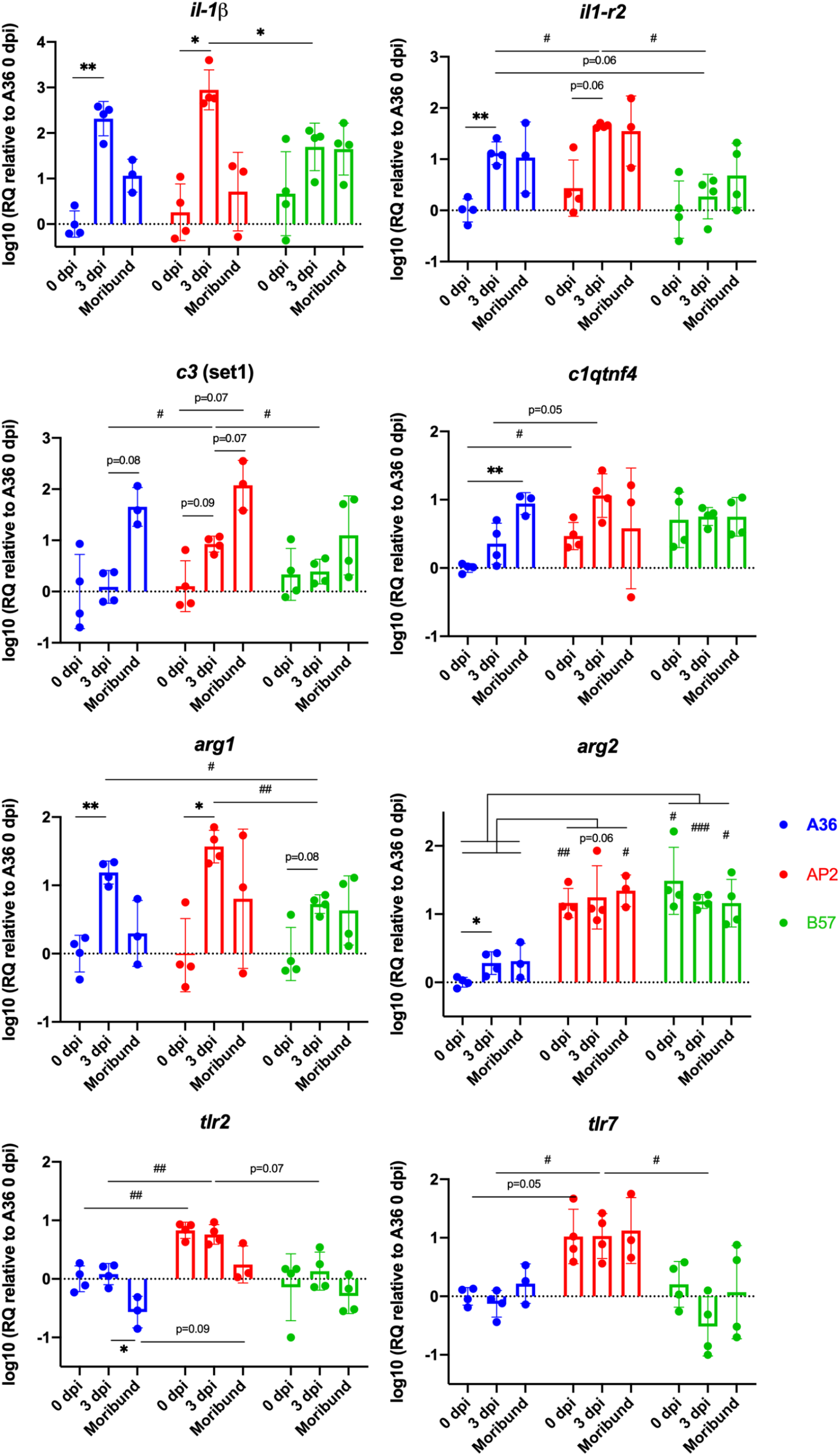
Relative quantification of gene expression in blood during *F. psychrophilum* infection of three isogenic lines. Gene expression is expressed as log10 (2^-∆∆Ct^) with A36 at 0 dpi as the reference condition. Moribund fish were sampled at 8, 11 and 14 dpi for A36, B57 and AP2, respectively. Values are the mean ± SD from 4 fish (except for AP2 and A36 which have only 3 surviving fish at the moribund datapoint). Values are compared using a mixed-effects model (REML) with GraphPad Prism 8.2.0 and differences between isogenic lines (#) or infection times (*) are considered significant with Bonferroni adjusted p-value < 0.0332 (*,#), <0.0021 (**,##) or <0.0002 (***,###). Representation of individual time-course expression values is available as Fig. S2.

The expression of the pro-inflammatory cytokine interleukin-1 beta (IL-1) in blood was significantly upregulated in isogenic lines A36 and AP2 at 3 dpi confirming a strong inflammatory response upon *F. psychrophilum* infection. Up-regulation was not significant in B57 due to a high level of expression before infection in 1 out of 4 fish. IL-1R2, a decoy receptor for IL-1 that is incapable of signaling and serves as negative regulator, and the C1q/TNF-related protein 4 (C1QTNF4), that acts as anti-inflammatory cytokine in macrophages [31], were also significantly up-regulated upon infection in lines A36 and AP2. IL-1R2 expression at 3 dpi was significantly higher in the Resistant line AP2 compared to the other ones. No significant change was observed in the expression of the anti-inflammatory cytokine IL-10 and of the multifunctional IL-11 upon *F. psychrophilum* infection in blood.

The complement component C3 plays a key role in antibacterial immunity by activating the complement cascade. Three functional forms named C3-1, C3-3 and C3-4 exist in rainbow trout and are the products of multiple paralogous genes [32] (Fig. S1). The C3 convertase cleavage site (Arg-Ser) is conserved in these three isoforms, but the factor I cleavage sites positions determining the binding specificity of the thioester-containing proteins are variable, potentially explaining different subfunctions [33]. In fact, up to eleven C3-like genes can be found in a recent high quality rainbow trout genome assembly supporting the hypothesis of a functional diversification. Several C3 encoding genes were found up-regulated upon *F. psychrophilum* infection in previous transcriptomic studies [24,25]. Here, we designed 3 sets of primers to target all gene copies encoding the functional C3-1, C3-3 and C3-4 forms as well as an additional uncharacterized C3 form (C3*) (Table 2; Fig. S1). C3-3 and C3-4 transcripts levels in blood were below the detection threshold but the C3-1 and C3* transcripts were quantified. Only C3-1 genes were up-regulated after infection in lines A36 and AP2 (Fig. 4; Fig. S2). For B57, a similar trend was observed for 2 out of 4 fish. Strikingly, the complement C3-1 genes were upregulated with a higher fold change and at early stage in the Resistant line AP2. The expression of other C3 genes did not change. These observations were consistent with the previous data reported in another BCWD Resistant isogenic line [24].

Expression of the arginase encoding genes *arg1* and *arg2*, two markers of M1/M2 polarized phenotypes of macrophages were also analyzed. It was noteworthy that *arg1* followed a typical pro-inflammatory pattern with upregulation in blood upon *F. psychrophilum* infection at 3 dpi in all isogenic lines. On the contrary, the expression of *arg2*, the marker of choice for M2 macrophages, which are associated with anti-inflammatory responses and wound healing [34], was not deregulated upon infection but strikingly, was expressed at significantly higher levels at steady state in lines AP2 and B57 compared to the Susceptible line A36.

The infection status did not modify the expression of *tlr2* and *tlr7* genes, as previously observed for TLR2, 3, 5, 9 and 22 in two other isogenic lines after *F. psychrophilum* infection [24]. However, the Resistant line AP2 showed a higher basal expression of *tlr2* and *tlr7* compared to the two other lines A36 and B57.

## 3. Discussion

Rainbow trout isogenic lines demonstrated variations in several traits such as responses to environmental stress factors including diet, temperature, confinement as well as susceptibility to different pathogens, indicating high genetic variability [20,35,36]. In an attempt to understand BCWD resistance traits, we identified and exploited three isogenic lines of rainbow trout with distinct susceptibilities to *F. psychrophilum*. The resistance/susceptibility status was not influenced by the route of infection for five isogenic lines challenged at an early life stage by bath (1 g) or injection (10 g), except for line B57 that tends to resist better by bath compared to injection. Strikingly, the relative resistance/susceptibility status was the same across life stages (from 1g to > 400g). Injection of as low as 10^2^ bacterial cells of strain FRGDSA 1882/11 produced high mortality in all isogenic lines at the juvenile stage, while a 100-fold higher infectious dose was required at the adult stage, exemplifying the typical susceptibility of young fish toward *F. psychrophilum* infection. When comparing the survival curves of each line from trials III and IV performed using the same infectious dose on adult fish with around 3-fold differences in body weight, only a slight delay in the onset of death events and lower mortality rate were observed when weight increased, indicating that fish size has a minimal impact on the BCWD resistance trait at the adult stage.

This study reports for the first time quantitative data on within-host growth of *F. psychrophilum* using blood sampling during the course of infection. The results confirm that BCWD is associated with massive septicemia in rainbow trout and that bacteremia appears as a good diagnostic indicator of *F. psychrophilum* infection. Moreover, we demonstrated that resistance is associated with a better control of pathogen growth. When natural protective skin barriers are overcome (such as mimicked here by injection), pathogen growth was nearly uncontrolled in Susceptible line A36, as exemplified by the generation time in blood (5.9 hours) that was only 2-fold longer compared to *in vitro* laboratory growth conditions (2.8 hours), and resulted in rapid death associated to high bacterial loads in organs. In contrast, bacterial load upon death were lower in lines B57 and AP2. Low dispersion of the bacterial load values was observed when death occurred, suggesting the existence of a lethal burden threshold that differs between isogenic lines. Strikingly, these values were comparable in B57 and AP2 while within-host growth rates and time-to-death significantly differed, indicating that these isogenic lines are both capable of restricting pathogen growth although more efficiently in AP2 than in B57. In addition, some AP2 survivors were able to support lethal burden, suggesting a putative tolerance mechanism in this BCWD Resistant line.

A better knowledge of BWCD pathogenesis will further contribute to developing diagnostic parameters to anticipate a prognosis and to identify host parameters for selective breeding. A study showed that the susceptibility of rainbow trout to *F. psychrophilum* infection is negatively correlated with spleen size [37]. Another study suggested a possible effect of relative frequencies of different myeloid cell subsets in spleen and head kidney [38]. The abundancy of neutrophil-like cells was negatively correlated to bacterial loads in infected organs suggesting that this specific cell type might support resistance to *F. psychrophilum* infection, in accordance with the great capacity of fish neutrophils to engulf surface associated microbes [39]. Based on our results, the pathogen dynamics in blood may also be a good indicator for predisposition of individuals to BCWD.

In an attempt to identify putative immune response differences between BWCD susceptible/resistant isogenic lines, we applied individual monitoring to analyze the basal level as well as expression changes upon infection on a selection of immune genes in blood. This 3Rs approach allowed us to minimize the number of animals for experimental infections. As previously reported for BCWD, signatures of pro-inflammatory responses were observed early during the infection process in all isogenic lines, as exemplified by the upregulation of IL-1 at 3 dpi.

BCWD resistance may arise from a stronger antibacterial activity. This is supported in this study by the early induction of complement C3 observed in blood in the Resistant line AP2 as well as by its higher expression previously observed in kidney for another Resistant line A03 (Langevin et al 2012). Our observations therefore point to the potential importance of C3-1 in the response against *F. psychrophilum*, which would deserve an in-depth characterization and correlation with genetic approaches.

Our data also suggest a role of arginase in BCWD. Arginine metabolism is a key regulatory mechanism of immune responses, as there is a competition for available intracellular arginine between arginase and nitric oxide synthase (NOS). Increase of arginase in the extracellular environment upon immune cell death also regulates arginine local concentration and modulates T cell function [40]. While *arg1* was upregulated in blood upon *F. psychrophilum* infection in all our isogenic lines, we observed no induction, but a higher expression of *arg2* at steady state in AP2 and B57 in comparison with A36. This expression pattern may contribute to the efficient resistance against *F. psychrophilum* infection. In common carp, Type II arginase was described as a probable marker for M2 macrophages whose role was found in anti-inflammatory action and tissue repair [34,41]. Besides, *arg2* expression was mainly detected in neutrophils in transgenic *arg2*:*GFP* zebrafish larvae early after tailfin transection, while very few M2 arg2+ macrophages were observed [42]. In this model, *arg2* was expressed both in neutrophil and macrophage subsets associated to granulomas after infection with *Mycobacterium marinum*. In our data, cell types constitutively expressing *arg2* in the blood remain unknown. Altogether, the correlation of this important anti-inflammatory factor which promotes early tissue regeneration with resistance to *F. psychrophilum* calls for further investigations. Indeed, a previous study comparing differences in tissue inflammatory damage between resistant and susceptible lines of rainbow trout found that a susceptible line was more likely to show a severe splenic necrosis lesions than a resistant one [43]. The higher expression of *arg2* in AP2 and B57 which were more resistant to *F. psychrophilum* infection could explain such pathologic difference.

TLRs are critical for the detection of pathogens and induction of primary responses. In mammals, they are the best-known pathogen associated molecular patterns (PAMPs) receptors which detect microbes and trigger rapid inflammatory responses, playing an essential role in the control of infections and initiation of adaptive immunity. The ligand specificities of TLRs in fish remain largely undetermined partially due to the low conservation of binding domains [44]. TLR2 is a membrane-bound receptor that supposedly binds lipopeptides of bacterial membranes. *tlr2* was shown to be induced by exposure to Gram-positive and Gram-negative bacteria in Indian major carp [45]; however, a study using fish cell lines failed to show its ability to induce the NF-ΚB pathway [46]. Until recently, molecular basis of LPS sensing in fish was not understood [44,47]. Higher basal expression of two TLR genes in the BCWD Resistant line AP2 compared to the Susceptible A36 and the Intermediate B57 lines may imply a better detection of PAMPs and more effective activation of antibacterial responses, as observed in AP2 with the earlier upregulation of complement C3 compared to the two other isogenic lines. High levels of antibacterial defenses in BCWD resistant fish are also consistent with the efficient limitation of bacterial growth observed within the host.

Focusing on leukocytes circulating in the blood, we encountered some differences compared to the previous study, for instance IL-10 anti-inflammatory cytokine expression was induced upon infection in the head kidney of B57 fish [24]. Similarly, the IL1-R2 showed higher basal expression in B57 in this tissue, but was not induced in the blood in our data. The difference in sampled materials likely explains these observations; for example, T cells are extremely scarce in rainbow trout blood, and other leukocyte subsets may reside in secondary lymphoid organs.

A similar immune gene expression kinetics study was conducted in susceptible and resistant BCWD rainbow trout lines [48]. Consistently with our observations, the pathogen load was significantly higher in the susceptible line. In addition, higher basal expression levels of putative cytokine receptors were found in spleen of the resistant fish. Together, the functional effects of these receptors in the resistance during the course of infection remain to be determined.

Our results indicate intrinsic variations in the response of the isogenic lines to *F. psychrophilum* infection. However, further investigation is required on whether this is related to differences in phagocytic efficiency or impedance to intracellular proliferation of *F. psychrophilum* among the isogenic lines. A study suggested *F. psychrophilum* as an intracellular bacterium by demonstrating increasing recovery of alive bacteria in phagocytes purified during the course of infection in rainbow trout [49]. A direct proof of proliferation inside host cells has not been reported yet, but would shed light on what immune mechanisms would be most efficient against this pathogen.

In summary, results from this study showed for the first time within-host bacterial growth and time-course innate immune responses using individual monitoring in blood of rainbow trout following infection with *F. psychrophilum*. The comparison of isogenic lines with contrasted susceptibilities to the disease revealed meaningful information regarding BCWD resistance traits which open promising perspectives for a better understanding of resistance mechanisms and identification of resistance predictors and genes.

## ACKNOWLEDGEMENTS

The authors are grateful to the staff of the fish facilities (IERP, INRAE, Infectiology of Fishes and Rodent Facility, doi: 10.15454/1.5572427140471238E12; and PEIMA, INRAE, Fish Farming Systems Experimental Facility, doi: 10.15454/1.5572329612068406E12) for supplying fish and for technical assistance and advice. This work was financially supported by the Agence Nationale de la Recherche (grant ANR-17-CE20-0020-01 FlavoPatho) and by institutional support from INRAE.

## AUTHOR CONTRIBUTIONS

Author contributions following the CRediT taxonomy (https://credit.niso.org) are as follows: Conceptualization: BL, EQ, PB, TR; Formal Analysis: BL, TR, PB; Funding acquisition: TR, EQ, ED; Investigation: BL, DR, ND, JFB, TR; Project administration: TR, PB; Supervision: TR, PB, ED; Writing – original draft: BL, TR; Writing – review & editing: BL, EQ, DR, ND, ED, JFB, PB, TR.

---

**Fig. S1.**
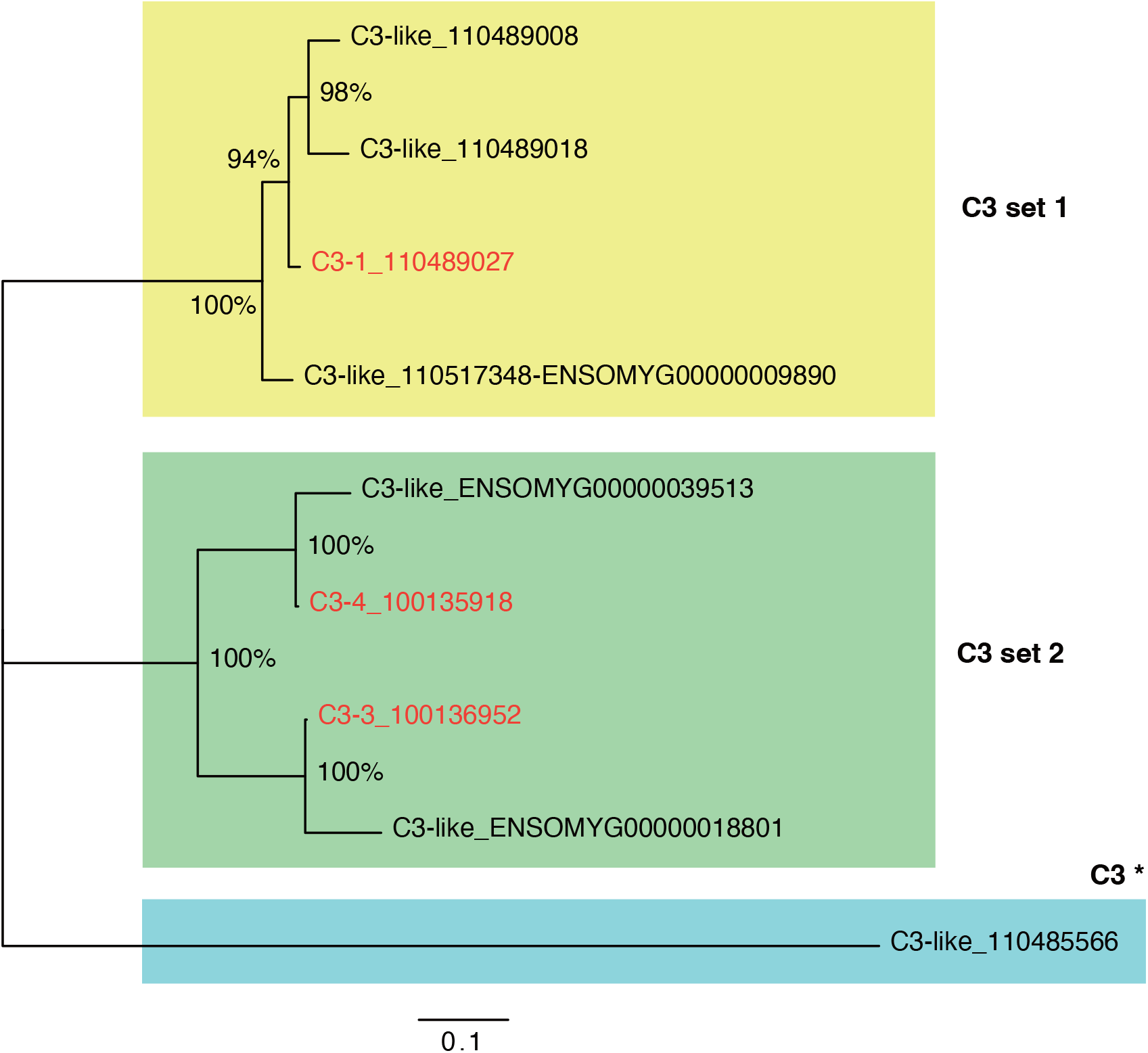
Unrooted phylogenetic tree of C3 protein sequences. C3-1, C3-3 and C3-4 sequences mentioned in the text are highlighted in red. Sequences amplified by our different primer sets (C3 set1, C3 set2, C3*) are boxed in yellow, green and blue. NCBI gene ID or Ensembl ENSOMYG ID are indicated for each tip. The evolutionary history was inferred by using the Maximum Likelihood method and the JTT matrix-based model. The tree with the highest log likelihood (−14308.73) is shown. The percentage of trees in which the associated taxa clustered together is shown next to the branches. Initial trees for the heuristic search were obtained automatically by applying the Neighbor-Join and BioNJ algorithms to a matrix of pairwise distances estimated using a JTT model, and then selecting the topology with superior log likelihood value. This analysis involved 9 amino acid sequences. There were a total of 1739 positions in the final dataset. Evolutionary analyses were conducted in MEGA X.

**Fig. S2.**
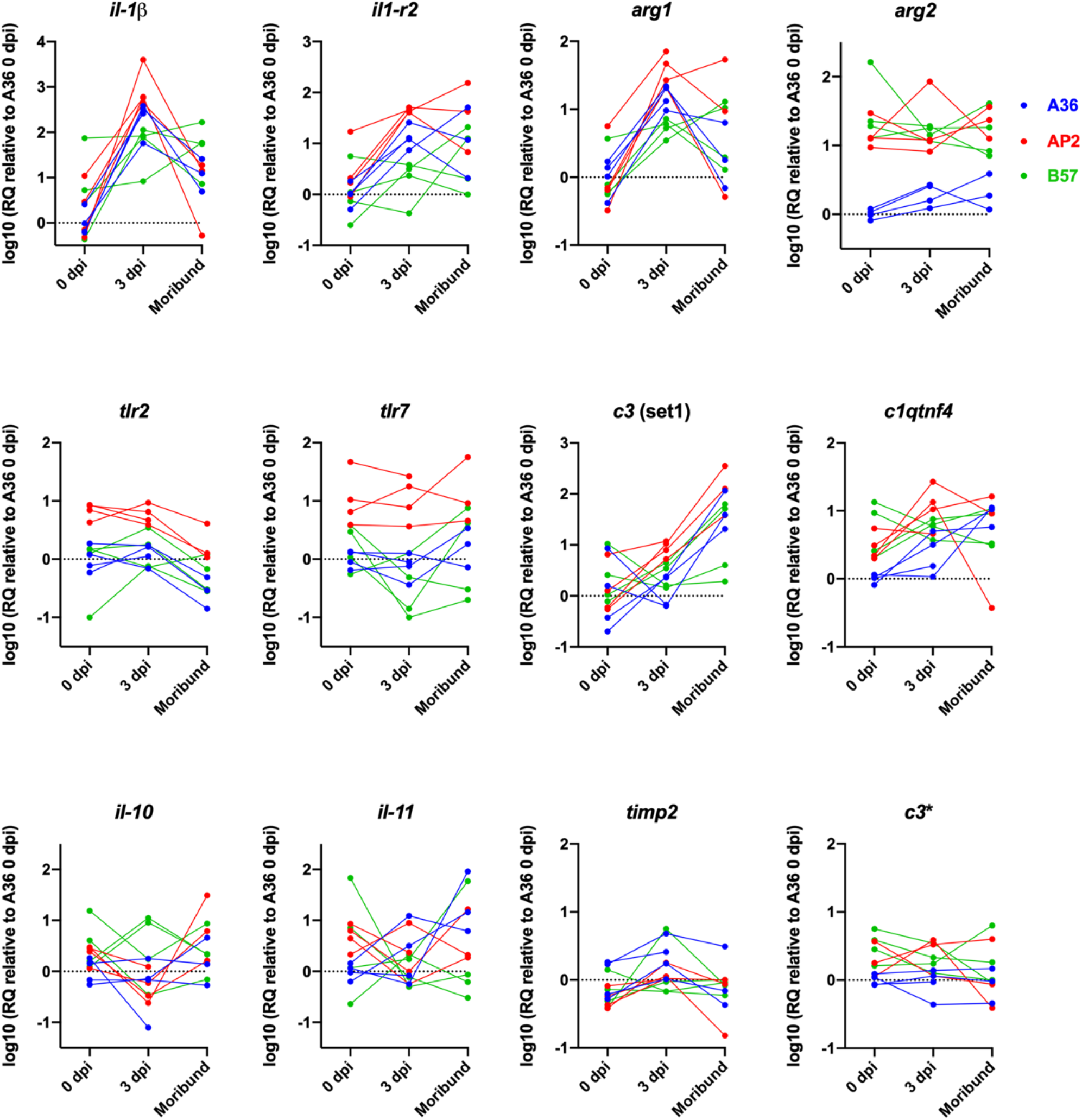
Relative quantification of gene expression at individual level in blood during *F. psychrophilum* infection of three isogenic lines. (same data as Fig. 4). Gene expression is expressed as log10 (2_-∆∆Ct_) with A36 at 0 dpi as the reference condition. Moribund fish were sampled at 8, 11 and 14 dpi for A36, B57 and AP2, respectively. Individual values are from 4 fish (except for AP2 and A36 which have only 3 surviving fish at the moribund datapoint).

## References

[1] FAO. The State of World Fisheries and Aquaculture 2020: Sustainability in action. Rome, Italy: FAO; 2020. https://doi.org/10.4060/ca9229en.

[2] Evensen Ø, Lorenzen E. An immunohistochemical study of Flexibacter psychrophilus infection in experimentally and naturally infected rainbow trout (Oncorhynchus mykiss) fry. Diseases of Aquatic Organisms 1996;25:53–61. https://doi.org/10.3354/dao025053.

[3] Madsen L, Dalsgaard I. Reproducible methods for experimental infection with Flavobacterium psychrophilum in rainbow trout Oncorhynchus mykiss. Dis Aquat Organ 1999;36:169–76. https://doi.org/10.3354/dao036169.

[4] Henriksen MMM, Madsen L, Dalsgaard I. Effect of hydrogen peroxide on immersion challenge of rainbow trout fry with Flavobacterium psychrophilum. PLoS One 2013;8:e62590. https://doi.org/10.1371/journal.pone.0062590.

[5] Fraslin C, Dechamp N, Bernard M, Krieg F, Hervet C, Guyomard R, et al. Quantitative trait loci for resistance to Flavobacterium psychrophilum in rainbow trout: effect of the mode of infection and evidence of epistatic interactions. Genet Sel Evol 2018;50:60. https://doi.org/10.1186/s12711-018-0431-9.

[6] Sundell K, Landor L, Nicolas P, Jørgensen J, Castillo D, Middelboe M, et al. Phenotypic and genetic predictors of pathogenicity and virulence in Flavobacterium psychrophilum. Front Microbiol 2019;10:1711. https://doi.org/10.3389/fmicb.2019.01711.

[7] Knupp C, Kiupel M, Brenden TO, Loch TP. Host-specific preference of some Flavobacterium psychrophilum multilocus sequence typing genotypes determines their ability to cause bacterial coldwater disease in coho salmon (Oncorhynchus kisutch). J Fish Dis 2021;44:521–31. https://doi.org/10.1111/jfd.13340.

[8] Henryon M, Berg P, Olesen NJ, Kjær TE, Slierendrecht WJ, Jokumsen A, et al. Selective breeding provides an approach to increase resistance of rainbow trout (Onchorhynchus mykiss) to the diseases, enteric redmouth disease, rainbow trout fry syndrome, and viral haemorrhagic septicaemia. Aquaculture 2005;250:621–36. https://doi.org/10.1016/j.aquaculture.2004.12.022.

[9] Silverstein JT, Vallejo RL, Palti Y, Leeds TD, Rexroad CE, Welch TJ, et al. Rainbow trout resistance to bacterial cold-water disease is moderately heritable and is not adversely correlated with growth. J Anim Sci 2009;87:860–7. https://doi.org/10.2527/jas.2008-1157.

[10] Fraslin C, Brard-Fudulea S, D’Ambrosio J, Bestin A, Charles M, Haffray P, et al. Rainbow trout resistance to bacterial cold water disease: two new quantitative trait loci identified after a natural disease outbreak on a French farm. Anim Genet 2019;50:293–7. https://doi.org/10.1111/age.12777.

[11] Leeds TD, Silverstein JT, Weber GM, Vallejo RL, Palti Y, Rexroad CE, et al. Response to selection for bacterial cold water disease resistance in rainbow trout. J Anim Sci 2010;88:1936–46. https://doi.org/10.2527/jas.2009-2538.

[12] Wiens GD, LaPatra SE, Welch TJ, Evenhuis JP, Rexroad CE, Leeds TD. On-farm performance of rainbow trout (Oncorhynchus mykiss) selectively bred for resistance to bacterial cold water disease: Effect of rearing environment on survival phenotype. Aquaculture 2013;388–391:128–36. https://doi.org/10.1016/j.aquaculture.2013.01.018.

[13] Wiens GD, Palti Y, Leeds TD. Three generations of selective breeding improved rainbow trout (Oncorhynchus mykiss) disease resistance against natural challenge with Flavobacterium psychrophilum during early life-stage rearing. Aquaculture 2018;497:414–21. https://doi.org/10.1016/j.aquaculture.2018.07.064.

[14] Wiens GD, Vallejo RL, Leeds TD, Palti Y, Hadidi S, Liu S, et al. Assessment of genetic correlation between bacterial cold water disease resistance and spleen index in a domesticated population of rainbow trout: identification of QTL on chromosome Omy19. PLoS One 2013;8:e75749. https://doi.org/10.1371/journal.pone.0075749.

[15] Vallejo RL, Palti Y, Liu S, Evenhuis JP, Gao G, Rexroad CE, et al. Detection of QTL in rainbow trout affecting survival when challenged with Flavobacterium psychrophilum. Mar Biotechnol (NY) 2014;16:349–60. https://doi.org/10.1007/s10126-013-9553-9.

[16] Vallejo RL, Liu S, Gao G, Fragomeni BO, Hernandez AG, Leeds TD, et al. Similar genetic architecture with shared and unique quantitative trait loci for bacterial cold water disease resistance in two rainbow trout breeding populations. Front Genet 2017;8. https://doi.org/10.3389/fgene.2017.00156.

[17] Palti Y, Vallejo RL, Gao G, Liu S, Hernandez AG, Rexroad CE, et al. Detection and validation of QTL affecting bacterial cold water disease resistance in rainbow trout using restriction-site associated DNA sequencing. PLoS One 2015;10:e0138435. https://doi.org/10.1371/journal.pone.0138435.

[18] Liu S, Martin KE, Gao G, Long R, Evenhuis JP, Leeds TD, et al. Identification of haplotypes associated with resistance to bacterial cold water disease in rainbow trout using whole-genome resequencing. Front Genet 2022;13:936806. https://doi.org/10.3389/fgene.2022.936806.

[19] Quillet E, Dorson M, Le Guillou S, Benmansour A, Boudinot P. Wide range of susceptibility to rhabdoviruses in homozygous clones of rainbow trout. Fish & Shellfish Immunology 2007;22:510–9. https://doi.org/10.1016/j.fsi.2006.07.002.

[20] Fraslin C, Quillet E, Rochat T, Dechamp N, Bernardet J-F, Collet B, et al. Combining multiple approaches and models to dissect the genetic architecture of resistance to infections in fish. Front Genet 2020;11:677. https://doi.org/10.3389/fgene.2020.00677.

[21] Pérez-Pascual D, Rochat T, Kerouault B, Gómez E, Neulat-Ripoll F, Henry C, et al. More than gliding: Involvement of GldD and GldG in the virulence of Flavobacterium psychrophilum. Front Microbiol 2017;8. https://doi.org/10.3389/fmicb.2017.02168.

[22] Barbier P, Rochat T, Mohammed HH, Wiens GD, Bernardet J-F, Halpern D, et al. The Type IX secretion system is required for virulence of the fish pathogen Flavobacterium psychrophilum. Appl Environ Microbiol 2020;86. https://doi.org/10.1128/AEM.00799-20.

[23] Zhu Y, Lechardeur D, Bernardet J-F, Kerouault B, Guérin C, Rigaudeau D, et al. Two functionally distinct heme/iron transport systems are virulence determinants of the fish pathogen Flavobacterium psychrophilum. Virulence 2022;13:1221–41. https://doi.org/10.1080/21505594.2022.2101197.

[24] Langevin C, Blanco M, Martin SAM, Jouneau L, Bernardet J-F, Houel A, et al. Transcriptional responses of resistant and susceptible fish clones to the bacterial pathogen Flavobacterium psychrophilum. PLOS ONE 2012;7:e39126. https://doi.org/10.1371/journal.pone.0039126.

[25] Marancik D, Gao G, Paneru B, Ma H, Hernandez AG, Salem M, et al. Whole-body transcriptome of selectively bred, resistant-, control-, and susceptible-line rainbow trout following experimental challenge with Flavobacterium psychrophilum. Front Genet 2015;5. https://doi.org/10.3389/fgene.2014.00453.

[26] Collet B, Urquhart K, Monte M, Collins C, Perez SG, Secombes CJ, et al. Individual monitoring of immune response in Atlantic salmon Salmo salar following experimental infection with infectious salmon anaemia virus (ISAV). PLOS ONE 2015;10:e0137767. https://doi.org/10.1371/journal.pone.0137767.

[27] Rochat T, Fujiwara-Nagata E, Calvez S, Dalsgaard I, Madsen L, Calteau A, et al. Genomic characterization of Flavobacterium psychrophilum serotypes and development of a multiplex PCR-based serotyping scheme. Front Microbiol 2017;8. https://doi.org/10.3389/fmicb.2017.01752.

[28] Duchaud E, Rochat T, Habib C, Barbier P, Loux V, Guérin C, et al. Genomic diversity and evolution of the fish pathogen Flavobacterium psychrophilum. Front Microbiol 2018;9:138. https://doi.org/10.3389/fmicb.2018.00138.

[29] D’Ambrosio J, Phocas F, Haffray P, Bestin A, Brard-Fudulea S, Poncet C, et al. Genome-wide estimates of genetic diversity, inbreeding and effective size of experimental and commercial rainbow trout lines undergoing selective breeding. Genetics Selection Evolution 2019;51:26. https://doi.org/10.1186/s12711-019-0468-4.

[30] Decostere A, Lammens M, Haesebrouck F. Difficulties in experimental infection studies with Flavobacterium psychrophilum in rainbow trout (Oncorhynchus mykiss) using immersion, oral and anal challenges. Res Vet Sci 2000;69:165–9. https://doi.org/10.1053/rvsc.2000.0408.

[31] Cao L, Tan W, Chen W, Huang H, He M, Li Q, et al. CTRP4 acts as an anti-inflammatory factor in macrophages and protects against endotoxic shock. Eur J Immunol 2021;51:380–92. https://doi.org/10.1002/eji.202048617.

[32] Sunyer JO, Zarkadis IK, Sahu A, Lambris JD. Multiple forms of complement C3 in trout that differ in binding to complement activators. Proc Natl Acad Sci U S A 1996;93:8546–51.

[33] Zarkadis IK, Sarrias MR, Sfyroera G, Sunyer JO, Lambris JD. Cloning and structure of three rainbow trout C3 molecules: a plausible explanation for their functional diversity. Dev Comp Immunol 2001;25:11–24. https://doi.org/10.1016/s0145-305x(00)00039-2.

[34] Wentzel AS, Petit J, van Veen WG, Fink IR, Scheer MH, Piazzon MC, et al. Transcriptome sequencing supports a conservation of macrophage polarization in fish. Sci Rep 2020;10:13470. https://doi.org/10.1038/s41598-020-70248-y.

[35] Dupont-Nivet M, Robert-Granié C, Le Guillou S, Tiquet F, Quillet E. Comparison of isogenic lines provides evidence that phenotypic plasticity is under genetic control in rainbow trout Oncorhynchus mykiss. J Fish Biol 2012;81:1754–62. https://doi.org/10.1111/j.1095-8649.2012.03437.x.

[36] Lallias D, Quillet E, Bégout M-L, Aupérin B, Khaw HL, Millot S, et al. Genetic variability of environmental sensitivity revealed by phenotypic variation in body weight and (its) correlations to physiological and behavioral traits. PLoS One 2017;12:e0189943. https://doi.org/10.1371/journal.pone.0189943.

[37] Hadidi S, Glenney GW, Welch TJ, Silverstein JT, Wiens GD. Spleen size predicts resistance of rainbow trout to Flavobacterium psychrophilum challenge. J Immunol 2008;180:4156–65. https://doi.org/10.4049/jimmunol.180.6.4156.

[38] Moore C, Hennessey E, Smith M, Epp L, Zwollo P. Innate immune cell signatures in a BCWD-resistant line of rainbow trout before and after in vivo challenge with Flavobacterium psychrophilum. Dev Comp Immunol 2019;90:47–54. https://doi.org/10.1016/j.dci.2018.08.018.

[39] Colucci-Guyon E, Tinevez J-Y, Renshaw SA, Herbomel P. Strategies of professional phagocytes in vivo: unlike macrophages, neutrophils engulf only surface-associated microbes. J Cell Sci 2011;124:3053–9. https://doi.org/10.1242/jcs.082792.

[40] Munder M, Schneider H, Luckner C, Giese T, Langhans C-D, Fuentes JM, et al. Suppression of T-cell functions by human granulocyte arginase. Blood 2006;108:1627–34. https://doi.org/10.1182/blood-2006-11-010389.

[41] Benedicenti O, Wang T, Wangkahart E, Milne DJ, Holland JW, Collins C, et al. Characterisation of arginase paralogues in salmonids and their modulation by immune stimulation/ infection. Fish Shellfish Immunol 2017;61:138–51. https://doi.org/10.1016/j.fsi.2016.12.024.

[42] Hammond FR, Lewis A, Anderson HE, Williams LG, Meijer AH, Wiegertjes GF, et al. An arginase 2 promoter transgenic illuminates anti-inflammatory signalling in zebrafish 2022:2022.02.14.480079. https://doi.org/10.1101/2022.02.14.480079.

[43] Marancik DP, Leeds TD, Wiens GD. Histopathologic changes in disease-resistant-line and disease-susceptible-line juvenile rainbow trout experimentally infected with Flavobacterium psychrophilum. Journal of Aquatic Animal Health 2014;26:181–9. https://doi.org/10.1080/08997659.2014.920735.

[44] Pietretti D, Wiegertjes GF. Ligand specificities of Toll-like receptors in fish: indications from infection studies. Dev Comp Immunol 2014;43:205–22. https://doi.org/10.1016/j.dci.2013.08.010.

[45] Samanta M, Swain B, Basu M, Panda P, Mohapatra GB, Sahoo BR, et al. Molecular characterization of toll-like receptor 2 (TLR2), analysis of its inductive expression and associated down-stream signaling molecules following ligands exposure and bacterial infection in the Indian major carp, rohu (Labeo rohita). Fish & Shellfish Immunology 2012;32:411–25. https://doi.org/10.1016/j.fsi.2011.11.029.

[46] Brietzke A, Arnemo M, Gjøen T, Rebl H, Korytář T, Goldammer T, et al. Structurally diverse genes encode Tlr2 in rainbow trout: The conserved receptor cannot be stimulated by classical ligands to activate NF-κB in vitro. Dev Comp Immunol 2016;54:75–88. https://doi.org/10.1016/j.dci.2015.08.012.

[47] Loes AN, Hinman MN, Farnsworth DR, Miller AC, Guillemin K, Harms MJ. Identification and characterization of zebrafish Tlr4 coreceptor Md-2. J Immunol 2021;206:1046–57. https://doi.org/10.4049/jimmunol.1901288.

[48] Kutyrev I, Cleveland B, Leeds T, Wiens GD. Proinflammatory cytokine and cytokine receptor gene expression kinetics following challenge with Flavobacterium psychrophilum in resistant and susceptible lines of rainbow trout (Oncorhynchus mykiss). Fish & Shellfish Immunology 2016;58:542–53. https://doi.org/10.1016/j.fsi.2016.09.053.

[49] Decostere A, D’Haese E, Lammens M, Nelis H, Haesebrouck F. In vivo study of phagocytosis, intracellular survival and multiplication of Flavobacterium psychrophilum in rainbow trout, Oncorhynchus mykiss (Walbaum), spleen phagocytes. Journal of Fish Diseases 2001;24:481–7. https://doi.org/10.1046/j.1365-2761.2001.00322.x.

